# Ecological inference and contaminant detection from fungal microbiome data with q2-fungal-traits

**DOI:** 10.64898/2026.06.17.732913

**Authors:** Anton Lavrinienko, Vinzent Risch, Chumei Tang, Annina Meyer, Lena Flörl, Nicholas A. Bokulich

## Abstract

Fungi are key members of microbial communities, yet microbiome surveys often lack trait-based information required for ecologically meaningful interpretation of mycobiome data. To demonstrate the value of fungal trait-based phenotyping in microbiome research, we re-analyzed N=3,221 samples across four case studies spanning human, agricultural, and environmental systems. In human cancer and vineyard datasets, trait-based analysis detected fungi producing macroscopic fruiting bodies, likely introduced via airborne spore dispersal, indicating widespread contributions of transient or contaminant fungi that can confound interpretation of sequencing data from tumor biopsies and grape berries. In sourdough fermentations, filamentous fungi were highly abundant alongside traditionally-recognized yeast and occupied distinct ecological niches. In forest soils, increasing habitat disturbance was associated with increased prevalence and abundance of plant pathogens, and a marked decline in ectomycorrhizal and lichenized fungi. These changes were accompanied by a shift toward large-spored taxa in urban soils, consistent with enhanced stress tolerance. To facilitate broader adoption of fungal phenotyping in microbiome studies, we introduce *q2-fungal-traits*, a QIIME2 plugin for automated integration of fungal taxonomy derived from marker-gene or shotgun metagenome sequencing surveys with ecological and functional trait data. The plugin assigns lifestyle-related traits and spore size estimates through hierarchical taxonomic matching and integrates directly into standard microbiome workflows. Our case studies demonstrate that integrating trait-based ecology with mycobiota datasets can generate novel findings and testable hypotheses, enabling inference of the functional (ir)relevance of community constituents. Our work contributes to bridging the gap between descriptive community profiling and functional ecology in microbiome research.

## Introduction

Microorganisms play fundamental roles in host-associated and environmental ecosystems, supporting essential functions and planetary health. In natural environments, they form complex communities comprising bacteria, fungi, archaea, viruses, and other microorganisms, collectively referred to as the microbiota. While advances in high-throughput DNA sequencing have revolutionized microbial ecology by enabling comprehensive surveys of microbial communities across diverse biomes, research efforts remain unevenly distributed. The bacterial component receives the vast majority of research attention, in favor of other key members of microbial ecosystems, including fungi. This imbalance is concerning because fungi are among the most taxonomically, phenotypically, and functionally diverse organisms on Earth [1], and contribute essential ecosystem services as well as important roles in pathogenesis across diverse environments, including plant and animal hosts [1, 2].

DNA sequencing enables untargeted taxonomic and/or functional profiling of complex microbial communities through marker-gene and whole metagenome sequencing approaches. For example, amplicon sequencing of the internal transcribed spacer (ITS) region of the ribosomal RNA locus has become the standard approach for cultivation-free profiling of fungal communities (mycobiota) [3, 4]. This strategy has been applied across diverse study systems (soil [5], air [6], animal gut [7, 8], and human cohorts [9, 10]), substantially expanding our understanding of fungal diversity and distribution. Although sequencing-based surveys overcome many challenges associated with fungal isolation and cultivation, the resulting datasets typically lack information on key fungal phenotypes and measurable traits, such as morphology and ecological lifestyle. The key issue is that the absence of such trait-based data limits our ability to interpret sequencing data patterns in a complete and ecologically meaningful manner.

The recent development and expansion of databases and community resources containing fungal functional trait information [11–15] provides an opportunity to close this gap by annotating sequencing-based taxonomic data with expert-curated information on fungal ecology and traits. Nevertheless, integration of trait-based knowledge into analyses of the mycobiota datasets remains relatively rare in the literature [15]. Indeed, the lack of routine cross-referencing between taxonomic and functional traits data in standard bioinformatics workflows likely reflects technical barriers to adoption and limited conceptual recognition of the potential benefits (and/or costs of ignoring fungal ecology) among researchers. At least partly, this is because available reference databases are not yet integrated into or fully interoperable with mainstream microbiome analysis workflows and often require manual curation or additional annotation steps outside the context of commonly used bioinformatics tools.

Here, we introduce *q2-fungal-traits*, a software package that enables automated trait-based classification of fungi from mycobiota profile data, including amplicon and shotgun metagenomic sequencing surveys. The *q2-fungal-traits* plugin links fungal taxonomy with expert-curated ecological and functional trait information available for thousands of fungal taxa, and can be integrated in a single step within the widely used QIIME 2 microbiome bioinformatics platform [16]. Using four case studies focused on fungi in (1) human cancer tissues, (2) vineyard ecosystems, (3) sourdough fermentations, and (4) forest soils, we demonstrate that integrating fungal trait information facilitates ecological interpretation and provides valuable, actionable insights about the fungal communities across diverse study systems and ecological contexts.

## Materials and Methods

### Automated trait-based classification of fungi

The QIIME 2 plugin *q2-fungal-traits* is a Python package that automates trait-based classification of fungi in microbiome sequencing data by linking fungal taxonomy with ecological and functional trait information, currently derived from two reference resources: (1) the FungalTraits, and (2) the fungal spore morphology databases. The FungalTraits database (v.1.2) contains expert-curated records for several lifestyle-related traits across 10,623 fungal and 142 fungus-like stramenopile genera [11]. This resource integrates information from other existing databases, including FUNGuild [17] and Fun^Fun^ [13], into a single comprehensive reference for trait-based annotation of fungal taxa [11]. The fungal spore size data were compiled by Aguilar-Trigueros et al. [6, 12], and include spore volume estimates for 25,789 fungal species. These data capture variation in the size of spores produced in both sexual (meiospores) and asexual (mitospores) cycles. Given that spore size varies considerably across fungal lineages and can influence dispersal potential and survival [12, 18], it represents an important yet often overlooked dimension of fungal ecology in mycobiome studies.

The method *qiime fungal-traits annotate* takes as input a taxonomy file containing hierarchical taxonomic annotations for fungal features, formatted according to commonly used reference taxonomy databases (UNITE [19] or NCBI [20] taxonomy). This method returns a tabulated file containing trait annotations generated through mapping user-provided taxonomic annotations to entries in the FungalTraits and spore morphology databases.

For traits derived from the FungalTraits, taxonomy strings are matched at the genus level, with additional validation against phylum-level identity to avoid incorrect matches among genera with divergent higher-level taxonomic lineages. Spore volume estimates are assigned using exact species-level matches for all features classified within the kingdom Fungi. Following the approach described by Abrego et al. [6], genus-level mean values, calculated based on the average spore volume estimate across all species assigned to a given fungal genus, are used when direct species-level estimates are missing. When genus-level data are also missing, family-level mean values are used to fill in annotation gaps. However, features lacking classification at the family level are not assigned spore volume estimates and are recorded as missing data. The resulting annotations include spore volume per taxon (in μm^3^), together with the taxonomic level used for each assignment (species, genus, or family).

*q2-fungal-traits* is implemented as a QIIME 2 plugin and integrates directly into standard QIIME 2 [16] and MOSHPIT [21] workflows for ITS amplicon and whole metagenome sequencing data analysis, supporting for instance filtering of features based on trait criteria, export of annotated data to common file formats, or visualization of taxonomy tables with associated trait annotations. The *q2-fungal-traits* is an open-source package, freely available together with usage examples at https://github.com/bokulich-lab/q2-fungal-traits.

### Trait and spore annotation coverage

To provide additional context and assess the coverage of fungal trait and spore size data, we evaluated the completeness of available annotations against the widely used UNITE reference taxonomy database [19] (release v.10, ITS sequences clustered at 99% identity, singletons excluded). Across 5,512 unique fungal genera in UNITE, FungalTraits coverage was highest for broad ecological and morphological descriptors, including primary lifestyle (78.9%), growth form (79.0%), fruiting body type (65.0%), and hymenium type (64.1%), whereas coverage varied across other traits (Figure 1 A). Among 27,639 unique fungal species in UNITE, more than 70% have associated spore size data, with most annotations assigned at the genus level (Figure 1 B). Coverage of spore size data was comparable for meiospores (85.0%) and mitospores (71.9%), whereas as expected, relatively few taxa (∼1.5%) had information on multinucleate spores [6, 12]. Our comparison of overlap between trait and spore annotations indicates that a substantial proportion of fungal species in UNITE are co-annotated with both data types. In particular, meiospore and mitospore size data were frequently available for taxa with lifestyle-and morphology-related trait annotations (Figure 1 C), suggesting that analyses focusing on these extensively annotated traits are likely to also benefit most from their combined use.

**Figure 1.**
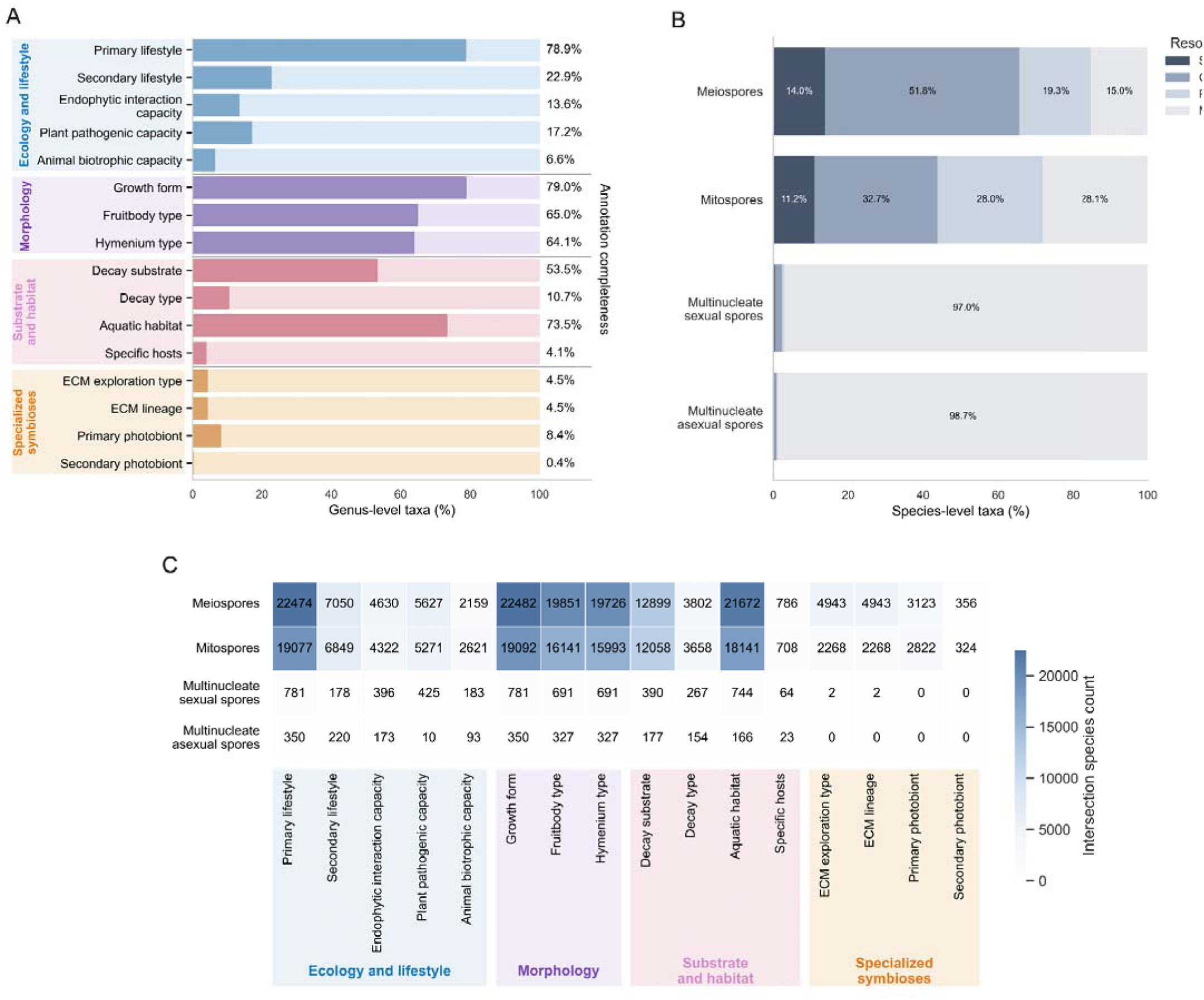
Coverage, taxonomic resolution, and overlap of fungal trait and spore size annotations. **(A)** FungalTraits annotation completeness across fungal genera (N=5,512) listed in the UNITE reference database, shown as the percentage of genera annotated for each trait-related field. **(B)** Spore volume annotation completeness and taxonomic resolution across fungal species (N=27,639) listed in the UNITE reference database, shown as the percentage of taxa assigned at species-, genus-, family-level, or missing for each spore type. **(C)** Intersection of fungal trait and spore size annotations across fungal species listed in UNITE. Each cell indicates the number of species carrying both annotation types; ECM, ectomycorrhizal. **Alt text:** *Multi-panel figure showing coverage of fungal trait and spore size annotations across taxa in the UNITE reference database. Bar plots summarize annotation completeness for fungal traits and spore size data, while a heatmap shows overlap between trait and spore annotations across fungal species, highlighting trait categories with the greatest shared coverage*.

### Case studies and data processing

We re-analyzed four publicly available mycobiota datasets. These case studies were selected to demonstrate how trait-based annotation of fungal taxa using *q2-fungal-traits* can facilitate ecological interpretation and provide additional insights into fungal communities across diverse study systems. Specifically, we used *q2-fungal-traits* to revisit the intrinsic mycobiota in (1) human cancer tissues and (2) vineyard ecosystems, and to examine broad ecological patterns in (3) sourdough fermentations, and (4) boreal forest soils.

In total, we processed ITS sequencing data from 3,221 samples across these four case studies, including (1) a total of 83,380,030 reads (mean 43,586; range 326-3,590,964) across N=1,913 tissue biopsy (1,584) and negative control samples (329, including 104 paraffin-only controls and 225 DNA-extraction controls) from Israeli multi-center cohort study investigating mycobiota signatures in distinct types of human cancer [22], (2) a total of 14,650,249 reads (mean 35,645; range 13-173,144) across N=496 grape berry (441) and soil (55) samples from Swiss vineyards [23], (3) a total of 11,839,187 reads (mean 23,726; range 6,392-50,283) across N=500 sourdough starters from a North American sourdough dataset [24], and (4) a total of 49,651,670 reads (mean 159,139; range 57,507-393,222) across N=312 samples from Finnish forest soils collected along a gradient of anthropogenic habitat disturbance [25].

Briefly, the raw ITS sequencing data for public mycobiota datasets were retrieved from the NCBI Sequence Read Archive using the *q2-fondue* plugin [26] in QIIME 2. The read data associated with samples from each of the four case studies were then processed using a standardized workflow in QIIME 2 [16]. Primer and adapter sequences were trimmed with the *cutadapt* plugin [27], after which forward reads were denoised and putative chimeric sequences removed using the *dada2* plugin [28] in QIIME 2. We then used *decontam* [29] through the *q2-quality-control* plugin to identify and remove potential contaminants based on their distribution in negative controls (where applicable). After these quality control steps, the resulting amplicon sequence variants (ASVs) were clustered into operational taxonomic units (OTUs) at 97% sequence similarity using the *cluster-features-closed-reference* method in the *q2-vsearch* plugin [30] against the UNITE reference database prepared with *RESCRIPt* [31] (UNITE release v.10, including ITS sequences for all eukaryotes clustered at 99% identity and excluding singletons [19]). Across all case studies, features not classified at the phylum level within the kingdom Fungi were removed prior to downstream analyses. The resulting feature table in the human cancer dataset (case study 1) included a total of 733 OTUs represented by 46,144,074 reads across all samples; data processing in the vineyard dataset (case study 2) resulted in 1,350 OTUs with a total of 10,367,639 reads in grape berry samples, and a total of 746 OTUs represented by 381,400 reads in soil samples; analysis of the sourdough dataset (case study 3) resulted in a total of 753 OTUs represented by 9,627,488 reads across all samples; and the forest soil dataset (case study 4) resulted in 5,354 OTUs represented by a total of 38,897,242 reads across all samples. These feature tables and corresponding taxonomic assignments based on the UNITE database were then used for trait-based annotation using the *q2-fungal-traits* plugin (https://github.com/bokulich-lab/q2-fungal-traits). To account for variation in sequencing depth among samples, in each case study feature tables were rarefied to an even sampling depth (i.e., (1) cancer dataset: 3,000 reads/sample, with a total of 3,459,000 reads and 702 OTUs across 1,153 remaining samples; (2) vineyard dataset: 10,000 reads/sample with a total of 3,370,000 reads and 1,276 OTUs across 337 remaining berry samples, and 5,000 reads/sample with a total of 200,000 reads and 705 OTUs across 40 remaining soil samples; (3) sourdough dataset: 4,500 reads/sample, with a total of 2,241,000 reads and 667 OTUs across 498 remaining samples; and (4) forest soil dataset: 37,266 reads/sample, with a total of 11,626,992 reads and 5,342 OTUs across 312 remaining samples), and unless otherwise stated, these normalized datasets were used for all downstream analyses.

We estimated mycobiota diversity by calculating several alpha diversity metrics, including richness (number of observed features), Shannon entropy, and evenness, for each sample using the *q2-diversity* plugin in QIIME 2. To assess the impact of trait-based feature filtering on alpha diversity estimates in our case studies, we compared alpha diversity between the original unfiltered sets containing all features and subsets containing features filtered either by (i) fruiting body type (e.g., ‘macrofungi’ [taxa producing macroscopic fruiting bodies], including taxa with agaricoid, apothecium (hymenium on surface), clathroid, clavarioid, corticioid, cyphelloid, gasteroid, gasteroid-hypogeous, hysterothecium (opening slith-like), mazaedium (pushpin-like), phalloid, polyporoid, and tremelloid fruiting body types), (ii) lifestyle-related traits (arbuscular mycorrhizal, ectomycorrhizal, and lichenized fungi [taxa forming complex multi-kingdom associations / specialized symbionts]), or (iii) fungal growth forms (e.g., yeast or fungi with filamentous mycelium) using Mann-Whitney U tests or paired Wilcoxon signed rank tests with Benjamini-Hochberg (BH) false discovery rate (FDR) correction for multiple comparisons. Note that in case study 1, we used a combination of (i) fruiting body type (macrofungi) and (ii) ecological lifestyle (specialized symbionts) traits to define ‘non-intrinsic’ fungal taxa and examine their distribution across human cancer mycobiomes. To examine differences in beta diversity between sample groups, we used the *q2-diversity* and *q2-kmerizer* plugins [32] in QIIME 2 to calculate distances between pairs of samples based on community composition (Jaccard index), as well as *k*-mer frequencies in considered features (*k*-mer based Jaccard index, captures subsequence-level information [32]). Dimensionality reduction was performed using principal coordinate analysis (PCoA) to visually examine sample clustering patterns, and significant differences among sample groups were tested with the permutational multivariate analysis of variance (PERMANOVA, 999 permutations) using *adonis* tests [33] implemented in the *q2-diversity* plugin in QIIME 2.

To examine community-wide shifts in spore size estimates between sample groups in the forest soil dataset, we calculated community weighted mean (CWM) spore volume per sample [34] either based on an average spore size of annotated taxa present in each sample (CWM weighted by presense-absence, with equal contribution across all taxa) or based on an average spore size of annotated taxa weighted by their relative abundance, thus taking into account dominance shifts in spore size across samples. The presence-absence weighted CWM and relative abundance weighted CWM were estimated for fungal meiospores, and differences in CWM spore sizes among study areas were then examined using Kruskal-Wallis tests with BH FDR correction.

Significant differences in trait prevalence and relative abundance distributions between sample groups were determined using Fisher’s exact tests and Kruskal-Wallis tests with BH FDR correction, respectively. To examine association between fungal growth forms and sample metadata in the sourdough dataset, we used a combination of correlation and variance-partitioning approaches. Feature tables were collapsed by growth form and transformed with the centered log-ratio (CLR) transformation to account for the compositional nature of the data. For numeric metadata variables (e.g., pH, latitude), CLR-transformed growth form abundances were correlated with metadata variables using Pearson correlations. For each growth form-metadata pair, we computed the squared correlation coefficient (R^2^) as an effect size and extracted associated *p*-values. For categorical metadata variables (e.g., country, flour type, container material), associations were assessed using analysis of variance (ANOVA), and eta-squared (η^2^) was calculated as a measure of effect size. To correct for multiple comparisons, p-values were adjusted per growth form using the BH FDR correction.

The code with all analysis steps and associated visualizations is available on GitHub: https://github.com/bokulich-publications/fungal-traits.

## Results

### Case study 1: human cancer tissue mycobiota

Cancer microbiome research is a rapidly growing field, driven by considerable interest in microbial signatures associated with different cancers and their potential diagnostic and predictive value [35–37]. A central challenge in this field is distinguishing intrinsic microbial diversity from non-target contaminants and host-derived signals in low-biomass tissue samples [38, 39]. Despite progress in establishing analytical and data reporting standards, the ecological relevance and intrinsic status of fungi detected in tumors remain relatively unexplored. In case study 1, we re-analyzed ITS data from 1,913 samples, including 1,584 tissue samples (i.e., tumor and normal adjacent tissue (NAT) from eight tissue types: breast, lung, melanoma, ovary, colon, brain, bone, and pancreas, as well as non-cancer normal breast tissue) and 329 negative controls from a multi-center study investigating mycobiota signatures across several major human cancer types [22].

We used *q2-fungal-traits* to examine the distribution of putative non-intrinsic fungal taxa across samples in the human cancer dataset based on their fruiting body types and lifestyle-related traits. We observed widespread detection of fungi that produce macroscopic fruiting bodies (macrofungi) both across various tissue types and in negative control samples (Figure 2 A-B). For example, these include edible mushrooms (*Agaricus bisporus, Collybia nuda, Pleurotus ostreatus*) as well as fungi associated with wood decay (*Coriolopsis gallica*, *Schizophyllum commune*, *Peniophora lycii*) (see examples: Figure 2 C), whose biology and macroscopic fruiting body type make colonization of human tissues highly improbable and which are thus unlikely to represent intrinsic components of cancer-associated fungal communities. Such non-intrinsic taxa were present in 4.5-58.8% of samples depending on sample type (Figure 2 A). Although most of these taxa occurred at low relative abundance, cumulatively non-intrinsic fungi often reached high abundance and in extreme cases accounted for more than 95% of total reads in individual samples (Figure 2 B). These patterns were consistent across medical centers, and were observed across all tissue types as well as in DNA-extraction negative controls and paraffin-only controls included in the original study, further suggesting a high probability that these fungal features represent environmental and/or reagent contaminants.

**Figure 2.**
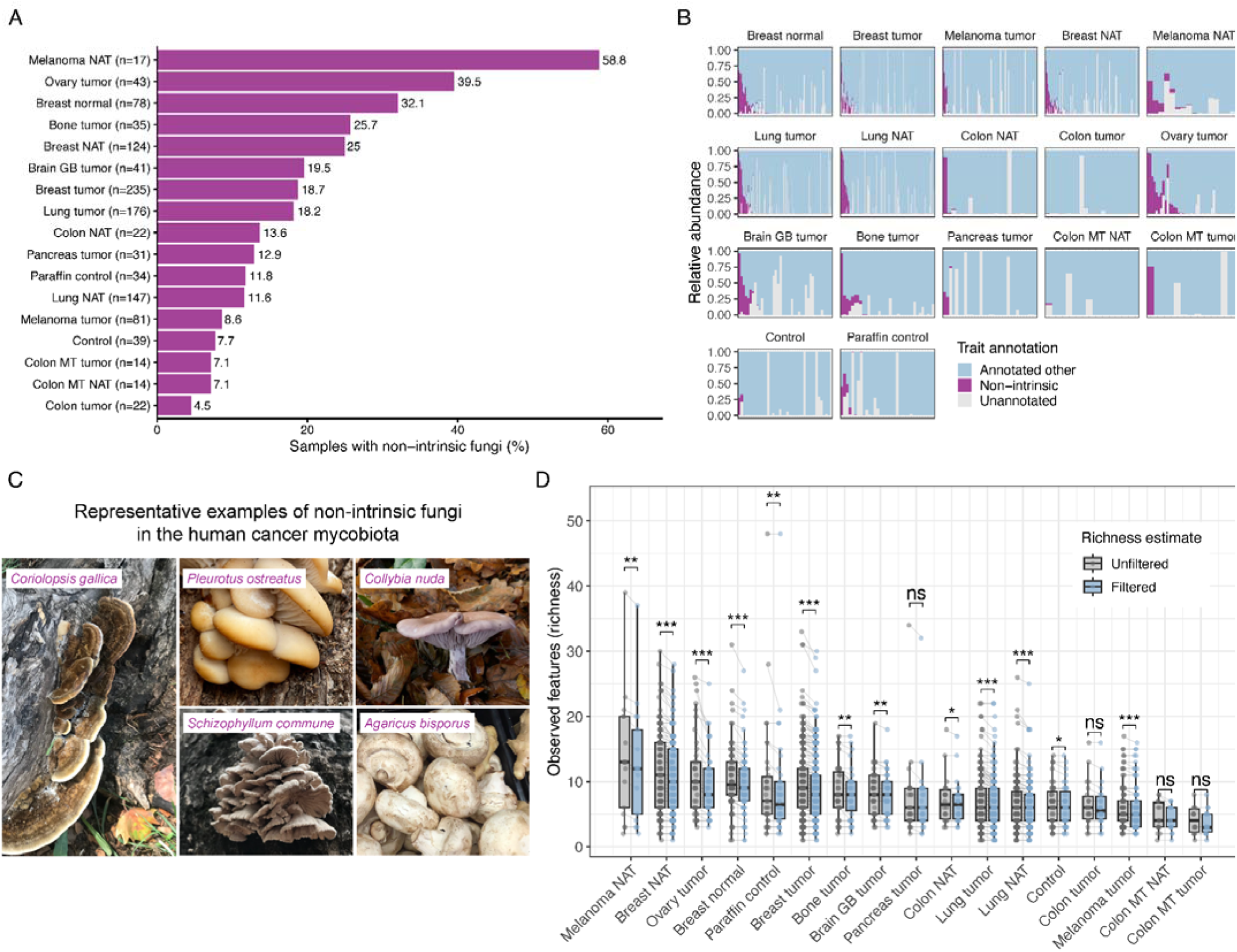
Distribution and impact of non-intrinsic fungal taxa across human cancer mycobiomes. **(A)** Show prevalence of non-intrinsic fungi across tissue types and negative control samples, displayed as the percentage of samples containing taxa classified as ‘non-intrinsic’ based on (i) fungal fruiting body type (‘macrofungi’) and (ii) ecological lifestyle-related trait annotations. Sample sizes for each sample type are indicated in parentheses; NAT, normal adjacent tissue; GB, glioblastoma; MT, metastatic. **(B)** Shows the mycobiota composition per sample, grouped by sample type. Bars represent the proportion of the community assigned to non-intrinsic fungi, annotated as other fungi, and taxa lacking trait annotation. Within each sample type, samples are ordered by decreasing relative abundance of non-intrinsic fungi. **(C)** Shows representative examples of non-intrinsic fungi in the human cancer mycobiota (within top 15 non-intrinsic features by prevalence and/or relative abundance across all samples). All images are CC0-licensed and sourced from iNaturalist.org. **(D)** Comparison of mycobiota richness before and after filtering non-intrinsic fungi. ‘Unfiltered’ richness estimates show the original dataset with all detected fungal features per sample, whereas ‘Filtered’ richness shows the subset that excludes features assigned to non-intrinsic fungi. Eac point represents a single sample and paired lines connect richness estimates before and after filtering within th same sample. Statistical significance was assessed using paired Wilcoxon signed rank tests, and all significanc indications are based on BH FDR corrected p-values, with p ≥ 0.05 = ns; * < 0.05; ** < 0.01; *** < 0.001. **Alt text:** *Multi-panel figure showing widespread detection of non-intrinsic fungi across human cancer mycobiomes. Macrofungi, including mushrooms and wood decaying fungi, are unexpectedly widespread across cancer types an negative control samples, and their removal substantially reduces estimated fungal richness*.

Although it is difficult to determine the exact impact of such non-intrinsic fungi on conclusions in the original pan-cancer analysis using this ITS amplicon sequencing dataset [22], their presence is likely to influence mycobiota diversity estimates and community composition patterns. Indeed, out of 702 unique fungal features detected across all samples, 110 (15.7%) were classified as non-intrinsic fungi. Our results also show that filtering these taxa significantly reduced alpha diversity estimates across most tissue types and negative control samples (Figure 2 D). Given the high relative abundance of non-intrinsic fungi in some samples, their inclusion may also affect downstream analyses, including those evaluating predictive value of identified taxa.

### Case study 2: vineyard mycobiota

Fungal communities play a central role in winegrowing as some taxa can shape wine characteristics [40]. Therefore, distinguishing between transient and resident taxa in grape-associated microbiomes is critical for the accurate interpretation of community assembly processes and fermentation dynamics. In case study 2, we re-analyzed ITS data from 441 grape berries and 55 soil samples originally collected across multiple Swiss vineyards to investigate environmental drivers of microbiome establishment in winegrowing ecosystems [23].

We annotated the retrieved OTUs with *q2-fungal-traits*, focusing on fruiting body type and specifically on the distribution of macrofungi. We found that the prevalence and abundance of macrofungi is highly dependent on the studied ecological niche. As expected, macrofungi were consistently detected in all soil samples and accounted for 21.07% of all OTUs, as this is a primary biome for hyphal growth and fruiting body formation. However, macrofungi were also highly prevalent on healthy grape berries, occurring in 96.14% of samples and representing 49.14% of all OTUs. Despite this high prevalence and diversity, most of these taxa occurred at low relative abundance (Figure 3 A-B). As a result, removing these taxa reduced the number of unique fungal features by nearly half, while decreasing total relative abundance by only 2.25%. Our findings indicate that many taxa recovered from berry samples are unlikely to represent resident members of the berry mycobiota and instead reflect transient inputs, likely from airborne spore deposition. What is important is that their inclusion substantially inflated alpha diversity estimates, significantly increasing community richness and Shannon entropy (Figure 3 C-E). This finding highlights the need to explicitly evaluate ecological relevance of taxa detected in mycobiome surveys and to make informed decisions about their inclusion in downstream analyses.

**Figure 3.**
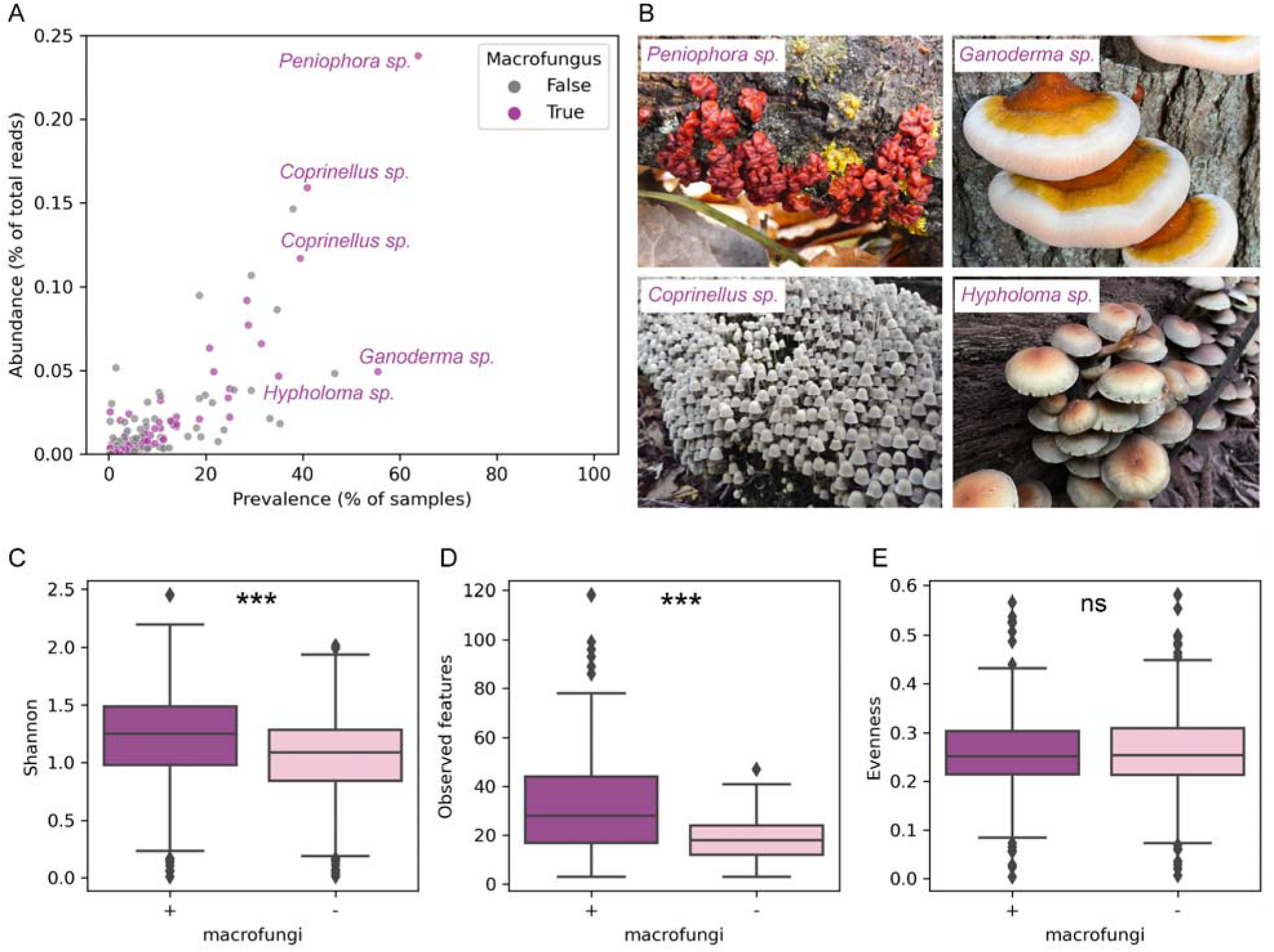
Distribution of fungi producing macroscopic fruiting bodies in the grape berry mycobiota. **(A)** Shows the prevalence and abundance of fungal OTUs in grape berry samples (abundance axis with cut off at 0.2 %), with features annotated as macrofungi highlighted in purple. **(B)** Shows representative examples of macrofungi in the grape berry mycobiota (within top 5 most prevalent and abundant features assigned to fungal genera annotate as macrofungi, i.e., *Peniophora*, *Ganoderma*, *Coprinellus*, *Hypholoma*). All images are CC0-licensed and source from iNaturalist.org. (**C–E**) Show differences in alpha diversity estimates between the original grape berry mycobiot dataset (+) and the resulting subset after filtering macrofungi (-), based on Shannon entropy (C), community richness (D), and evenness (E). Statistical significance was assessed using Mann-Whitney U tests, with significance indications based on p ≥ 0.05 = ns; * < 0.05; ** < 0.01; *** < 0.001. **Alt text:** *Multi-panel figure showing the prevalence, abundance, and diversity impacts of fungi producing macroscopic fruiting bodies in grape berry mycobiomes. Macrofungi are widespread despite their low relative abundance in the grape berry mycobiota, and their removal significantly reduces estimates of fungal community diversity*.

Analysis of lifestyle-related traits further confirmed clear ecological differentiation between sample types. Berry-associated communities were dominated by sooty molds (68.58% relative abundance), putative plant pathogens (21.98%), and nectar/tap saprotrophic fungi (5.82%), consistent with a sugar-rich, plant-associated environment. In contrast, soil communities exhibited greater functional diversity, including similar proportions of plant pathogens (21.27%) but also substantial representation of soil, litter, and wood saprotrophs. Notably, a large fraction of soil taxa remained unassigned (∼40% of relative abundance), likely reflecting both the higher diversity of soil fungal communities and current limitations in trait annotation coverage.

### Case study 3: sourdough mycobiota

Sourdough is a back-slopped (serially propagated) mixture of flour and water that develops through fermentation into a stable microbiome that determines sourdough bread quality [41]. Yeast are typically considered the dominant functional group of fungi in sourdough starters, primarily due to their facultatively fermentative metabolism responsible for carbon dioxide production for leavening and flavor formation. However, the presence and relative contribution of fungi with other growth forms remains underexplored. In case study 3, we re-analyzed ITS data from 500 sourdough samples originally collected to investigate microbial interactions and biogeographic patterns in sourdough starter communities in North America [24].

We annotated OTUs retrieved from sourdough samples using *q2-fungal-traits*, focusing on annotations of fungal growth forms. Contrary to the conventional yeast-centric model of sourdough fermentation, we found that filamentous mycelium-forming fungi were both prevalent and abundant across samples (Figure 4 A), accounting for 18% of total reads and nearly twice as many unique features as yeast-like fungi. In some sourdough starters, filamentous fungi even exceeded yeast in relative abundance (Figure 4 B), indicating that sourdough mycobiomes are not universally dominated by yeast.

**Figure 4.**
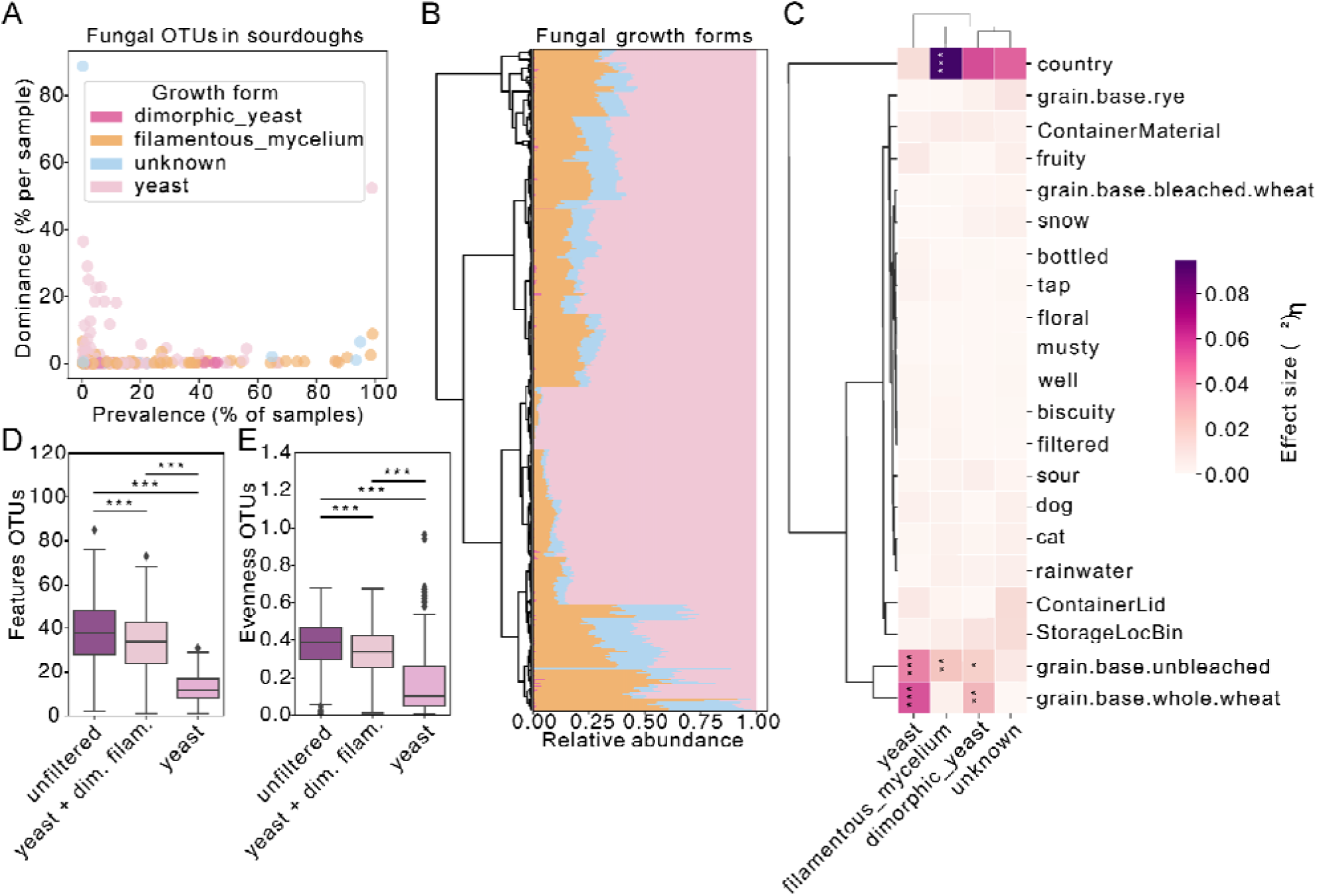
Fungal growth forms in the sourdough mycobiota. **(A)** Prevalence-dominance plot showing that filamentous fungi constitute a highly prevalent component of sourdough mycobiota. Each point represents a single fungal OTU. **(B)** Shows the composition of fungal growth forms per sourdough sample, indicating that filamentous fungi often rival or even exceed yeast in relative abundance. **(C)** Anova-based explained variance analysis across growth forms and sample metadata shows that filamentous fungi vary widely between countries, while yeast abundance is more variable across flour substrates (e.g., wholemeal, bleached/unbleached). (**D** and **E**) Show that restricting mycobiota analysis to yeast-only OTUs results in strong reduction in community richness (D) and evenness (E); statistical significance was assessed by Mann-Whitney U tests. All significance indications are based on BH FDR corrected p-values, with p ≥ 0.05 = ns; * < 0.05; ** < 0.01; *** < 0.001. **Alt text:** *Multi-panel figure showing the distribution and ecological importance of different fungal growth forms in sourdough mycobiomes. In addition to yeast, filamentous fungi are a common and abundant component of sourdough communities. They show different ecological associations than yeast and contribute substantially to overall fungal diversity*.

Our analysis of trait-metadata associations revealed distinct ecological patterns linked to fungal growth forms in sourdough mycobiomes. For example, filamentous fungi varied strongly with geography, whereas yeast abundance was more consistently associated with flour type, including differences between wholemeal and refined flours (Figure 4 C). Strikingly, limiting analyses to yeast-only taxa resulted in the loss of more than half of all fungal features (Figure 4 D) and also led to marked reduction in community evenness (Figure 4 E). As such, our results demonstrate that excluding non-yeast taxa, as is commonly done in published literature [24], can substantially skew ecological interpretation and underestimate fungal diversity in sourdough starters.

Notably, a fraction of the dataset (8.3% of reads representing 132 OTUs) remained unassigned with respect to growth form. These taxa, primarily within Ascomycota families such as *Nectriaceae* and *Pleosporaceae*, reflect current limitations of trait-based inference approach and taxonomic resolution in ITS-based short-read amplicon sequencing surveys.

### Case study 4: forest soil mycobiota

Urbanization represents one of the fastest-growing forms of environmental change globally and provides a powerful natural experiment for studying microbiome responses to anthropogenic disturbance. Although several studies have documented shifts in both environmental [25] and host-associated [42, 43] microbiomes between natural and urbanized habitats, most available studies remain largely descriptive and provide limited mechanistic insight into compositional turnover in microbiomes across disturbed environments. In the case study 4, we re-analyzed ITS data from 312 soil samples from natural forests located in national parks (N=105), managed forests (N=112), as well as suburban (N=48), and urban forests (N=47), originally collected to investigate the impacts of anthropogenic disturbance on the forest soil microbiome [25].

We used *q2-fungal-traits* to examine the distribution of lifestyle-related traits and spore volume estimates in soil fungi across a gradient of habitat disturbance. We found clear differences in the forest soil mycobiota composition between study areas (Figure 5). Specifically, increasing habitat disturbance was associated with a significant increase in the prevalence and/or relative abundance of fungal genera assigned to plant pathogens and animal parasites, accompanied by a marked decline in ectomycorrhizal and lichenized fungi (Figure 5 A-B). Notably, the largest differences in lifestyle-related traits were consistently observed between fungal communities in urban forests and national parks, indicating strong functional turnover at opposite ends of the habitat disturbance gradient. More broadly, these patterns suggest a transition from communities dominated by mutualistic taxa toward those characterized by pathogenic and opportunistic lifestyles under increasing anthropogenic disturbance.

**Figure 5.**
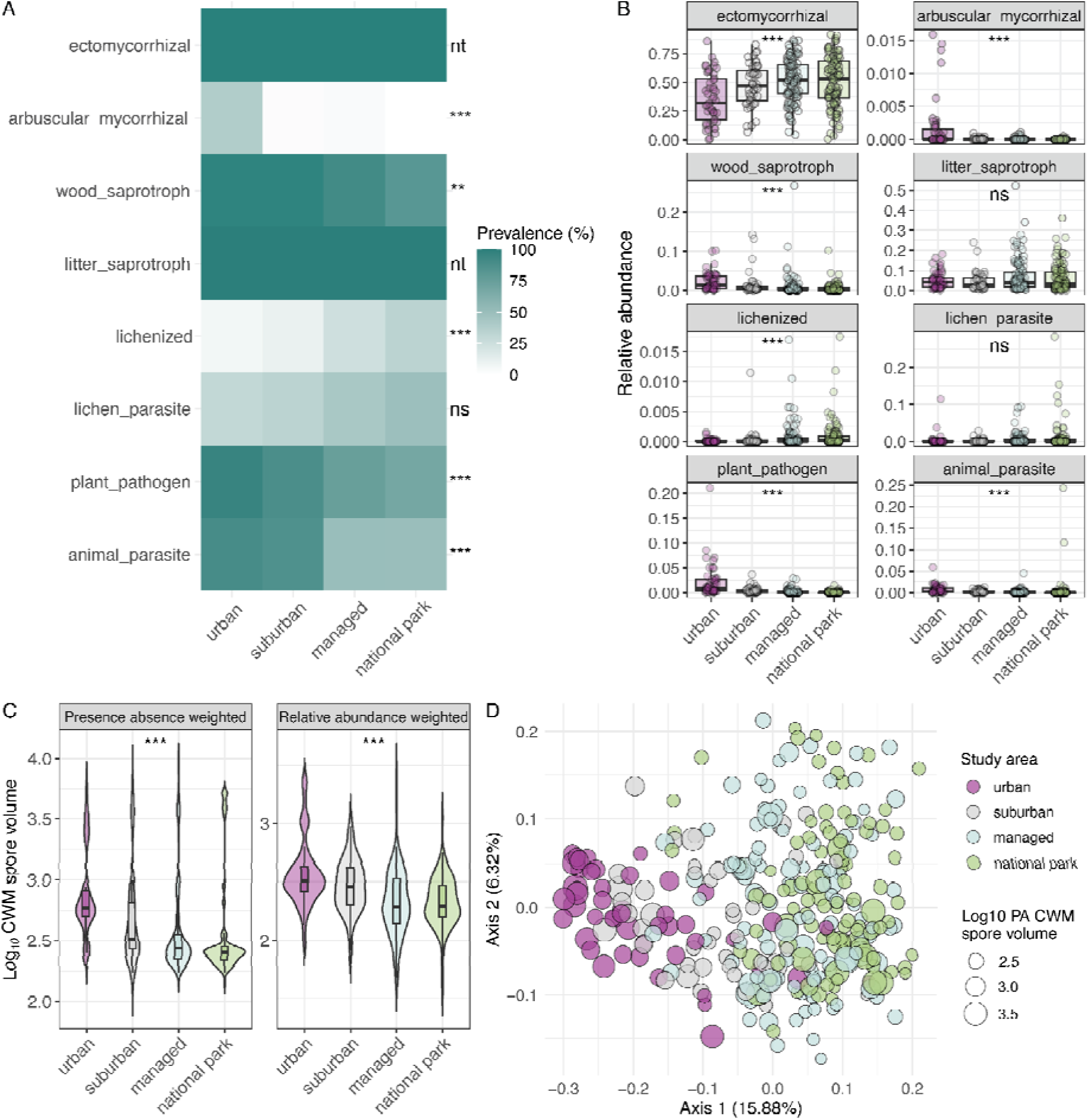
Distribution of lifestyle-related traits and spore size estimates in forest soil fungi. **(A)** Shows the prevalence and **(B)** relative abundance of major fungal lifestyle traits in soil samples collected from study areas alon a gradient of habitat disturbance (from urbanized to natural forest habitats). **(C)** Shows differences in presence-absence (PA) based and relative abundance (RA) based community weighted mean (CWM) meiospore volume between study areas. **(D)** Shows PA-based CWM meiospore volume data in relation to the soil mycobiota bet diversity (kmer-based Jaccard index) patterns. Statistical significance was assessed using Fisher’s exact tests for trait prevalence data (A), and Kruskal-Wallis tests for trait relative abundance (B) and spore size (C) data. Significant differences in beta diversity between study areas (D) were determined using permutation-based statistical tests (PERMANOVA, R^2^=0.115, p < 0.001). All significance indications are based on BH FDR corrected p-values, with p 0.05 = ns; * < 0.05; ** < 0.01; *** < 0.001; in panel (A), nt = not tested because prevalence did not vary among samples. **Alt text:** *Multi-panel figure showing changes in fungal lifestyle traits and spore sizes in forest soils across a gradient of habitat disturbance. Forest soil mycobiota in urban areas contain fewer ectomycorrhizal and lichenized fungi, more plant pathogens, and larger average spore sizes compared to natural forest habitats*.

Trait-based analysis of dispersal related characteristics also revealed concurrent shifts in spore size distributions. Community weighted mean (CWM) spore volume increased with urbanization, with significant shifts observed both when CWM was calculated using presence-absence weighting and when weighted by taxa relative abundance (Figure 5 C-D). This consistent pattern likely reflects combined effects of taxonomic turnover and changes in dominance structure, suggesting that taxa that produce larger spores become more prevalent and more abundant in urban soils. These observations may have direct functional relevance given that differences in fungal spores size can influence their dispersal and germination potential [12, 18, 34, 44, 45]. Together, our results provide a mechanistic hypothesis into how soil fungi respond to urbanization which can be tested in future studies.

## Discussion

Integrating phenotypic trait information describing morphology and lifestyle traits provides a critical link between taxonomic profiling and ecological function in large-scale microbiome surveys. Without this information, sequencing-based analyses remain largely descriptive, limiting our ability to interpret community patterns mechanistically and within an ecologically relevant context. Although specialized trait databases and tools exist (e.g., FungalTraits [11], FUNGuild [17], Fun^Fun^ [13]), their integration is relatively rare in routine microbiome analyses [15]. By embedding trait annotation directly within QIIME 2 [16], *q2-fungal-traits* lowers barrier in their adoption and allows for seamless and reproducible integration with widely used bioinformatics pipelines and downstream diversity analyses. We anticipate that this standardization will promote broader use of trait-based approaches in mycobiome studies.

Novel results from our case studies highlight the importance of explicitly considering the ecological origin of detected taxa in sequencing-based mycobiome studies. In analyses of human cancer [22] and vineyard ecosystems [23], we frequently detected fungi producing macroscopic fruiting bodies (macrofungi) and other taxa whose ecological traits make colonization of these environments highly unlikely. For example, in human tissue samples several taxa such as edible and cultivated mushrooms were consistently identified across cancer types and study centers, and also in DNA-extraction blanks and paraffin-only control samples [22]. Although their exact source remains unclear, many of these taxa are globally distributed and produce abundant airborne spores, suggesting dispersal from environmental sources to sample processing facilities in study hospitals and/or laboratories. It could be interesting to further test such putative dispersal routes, for instance, by examining mycobiota profiles from matched tumor tissues processed in conventional vs. specialized clean facilities [39]. Similarly, although fungal fruiting bodies are not expected on healthy grapes, soils from the same vineyards are enriched in such taxa and hence likely act as a source of spores detected on grape berries. Interestingly, taxa assigned to the genus *Peniophora* were consistently the most prevalent and abundant macrofungi in both human cancer and vineyard mycobiota case studies. This pattern across independent datasets is consistent with the ecology of *Peniophora*, a cosmopolitan genus of saprotrophic wood decay fungi that produce small spores with high potential for long-distance airborne dispersal [46]. Further evidence supporting an aerial origin of these taxa in our case studies comes from the Global Spore Sampling Project, showing that plant pathogens and wood saprotrophs represent some of the most abundant fungal groups detected through air sampling [6]. Regardless of their origin, in both of these case study examples, such taxa are unlikely to colonize these niches and most likely represent environmental contaminants that will not play significant roles in host health or wine quality. Together, our findings demonstrate that a substantial fraction of sequencing-based signals may reflect transient and/or contaminant inputs rather than intrinsic or resident members of fungal communities.

Our data clearly show that in addition to extensive quality control steps and filtering of host-derived reads in cancer mycobiome studies [35, 39, 47, 48], it is critical to assess the ecological relevance of detected fungi. The substantial inflation of alpha diversity caused by contaminant fungal taxa in our cancer mycobiota case study demonstrates that overlooking this issue can significantly distort mycobiota diversity estimates, which remain among the most commonly reported outcomes in the microbiome literature. Moreover, as the microbiome field increasingly moves toward predictive and translational applications, the consequences of ignoring fungal ecology likely extend beyond diversity estimates. For example, inclusion of non-intrinsic fungal taxa and other biological false-positive microbial signals, could generate spurious associations and influence the discovery of microbiome-based diagnostic and/or prognostic biomarkers. This issue is particularly relevant in cancer microbiome research, where identifying predictive microbial signatures is often one of the central research objectives [22, 36–38, 48].

An important question is how such signals should be handled in microbiome studies. Existing guidelines address contamination risks and focus on best practices from experimental design to molecular and bioinformatics analyses [38, 39, 49–51], including recommendations specific to mycobiome studies [4, 10, 52]. However, the issue of non-intrinsic taxa we demonstrate here differs from typical sources of technical noise or contamination described in the literature. Rather than uniformly filtering taxa with ecologically unexpected traits, we advocate for informed and context-dependent evaluation of their potential ecological and functional relevance based on sample type, study system, and experimental design [49, 53]. We recognize that trait annotations alone may not always provide definitive evidence for functional relevance (or irrelevance) of community constituents, yet they can offer valuable ecological context that can help distinguish intrinsic community members from transient taxa or potential contaminants. For example, in our vineyard mycobiota case study, fungi producing macroscopic fruiting bodies may be intrinsic to and functionally relevant members of soil communities but not to grape berry mycobiomes, where they are more likely to reflect transient spore contaminants. On the other hand, even such environmental signatures could be valuable in some contexts, e.g., for microbial source tracking or forensics applications. Again, careful evaluation of such signals is likely to be particularly important in cancer mycobiome studies and other study systems with low microbial biomass samples, where contamination can disproportionately influence results [39, 49]. Trait-based annotation offers a practical and scalable solution to address this issue and mitigate at least some false-positive microbial signals by routinely screening ecological characteristics of detected taxa, supporting more accurate interpretation of mycobiota diversity and composition.

Trait-based annotation can also power a shift from purely taxonomic descriptions toward ecosystem-level interpretation of fungal communities, as exemplified by other findings in our case studies. In sourdough fermentations [24], growth form based analyses revealed consistent differences between filamentous fungi and yeast, suggesting that these two fungal groups respond to distinct ecological cues (biogeography vs. substrate type) and occupy complementary niches within sourdough communities. In forest soils [25], trait-based profiles captured a transition from mutualistic to pathogenic functional groups along an urbanization gradient, accompanied by apparent shifts in dispersal related traits. The increase in large-spored taxa in areas with greater anthropogenic disturbance may have ecosystem-level functional consequences [54]. While fungi with larger spores are expected to disperse over shorter distances, they are generally considered more stress resistant and better adapted to survive and germinate under variable environmental conditions [12, 18, 34, 44, 45], which could be particularly useful in the context of habitat disturbance in urban areas. Interestingly, similar patterns were reported in a recent independent study comparing fungal communities in soils from urban and natural areas, suggesting that these observations represent consistent and generalizable ecological trends [34, 55]. These examples demonstrate how integrating functional traits into the sequencing-based mycobiome surveys can reveal ecological patterns that are not readily apparent from taxonomic profiles and compositional data alone, and support a more mechanistic understanding of fungal community dynamics across diverse study systems and ecological contexts [18, 54].

Despite these advances, several limitations remain. Methodological factors can influence trait-based ecological inference. For example, primer choice and DNA extraction protocols can influence the detection of specific fungal groups and associated traits. Indeed, detection of fungi producing rigid, hard-to-lyse spores may depend on lysis efficiency during DNA extraction. In addition, the ITS marker region commonly used for mycobiota profiling has known limitations for certain ecologically important groups. For example, arbuscular mycorrhizal fungi often require alternative markers because ITS variability limits reliable amplification and taxonomic resolution in this lineage using some primer sets [56, 57]. As a result, trait-based analyses relying on ITS data may underestimate specific functional groups, and complementary markers or targeted approaches may be required to fully capture fungal functional diversity. These technical factors should be considered when interpreting trait-based patterns from fungal sequencing data.

Our analyses show that a substantial proportion of fungal diversity in reference databases such as UNITE [19] lacks trait annotation, resulting in incomplete and uneven coverage across taxonomic groups. This limitation reflects both the diversity of fungi and current gaps in our understanding of fungal ecology, highlighting the need for continued efforts and investments in database expansion. Recent advances in machine learning and artificial intelligence offer opportunities to accelerate this process. For example, automated mining of phenotypic and ecological information from databases, culture collections, and specialized literature (e.g., MycoBank [https://www.mycobank.org/], resources maintained by the Westerdijk Fungal Biodiversity Institute [https://wi.knaw.nl/], MycoKeys publications), could facilitate scalable extraction and standardization of fungal trait data. Expanding trait databases to improve taxonomic coverage and incorporate additional dimensions, including for instance genomic traits such as rDNA copy number variation [58, 59] and metabolic characteristics analogous to BacDive ([60], https://bacdive.dsmz.de/), would improve analytical resolution and broaden trait-based applications. Advancing these resources will strengthen the integration of fungi into predictive microbiome research and overall support more mechanistic and scalable trait-based microbial ecology. As such resources become available, we would plan to upgrade *q2-fungal-traits* and invite contributions from the mycobiome community to enhance the capabilities of this plugin for new trait-based inference.

Taken together, our case studies demonstrate that integrating fungal trait information into microbiome analyses enables more ecologically meaningful interpretation of sequencing data, which can generate novel findings and testable hypotheses about community assembly and function across diverse study systems. By automating the integration of fungal taxonomy with ecological and functional traits within standard QIIME 2 workflows, *q2-fungal-traits* lowers technical barriers and facilitates scalable trait-based analyses in mycobiome studies. As such, our work highlights the potential of trait-informed microbiome analyses and provides a practical approach to bridge the gap between descriptive community profiling and functional ecology in microbiome research.

## Data Availability Statement

All analyses presented in this manuscript are based on publicly available datasets accessible through the NCBI Sequence Read Archive (SRA). The datasets associated with the four case studies are available under the following BioProject accession numbers: (1) human cancer mycobiota [22], accession PRJNA786764; (2) vineyard mycobiota [23], accession PRJEB89112; (3) sourdough mycobiota [24], accession PRJNA589612; and (4) forest soil mycobiota [25], accession PRJNA823643. The code with all analysis steps and associated visualizations presented here is available on GitHub: https://github.com/bokulich-publications/fungal-traits.

The fungal trait and spore size reference data implemented in the *q2-fungal-traits* plugin are available through the original associated publications [6, 11, 12]. The *q2-fungal-traits* is an open-source package, freely available following the installation instructions at https://github.com/bokulich-lab/q2-fungal-traits.

## Acknowledgements

We would like to thank Reinhard Berndt, curator of the Fungarium at ETH Zurich, for helpful discussions on the classification of macrofungi using fungal traits. We also thank Sven Stoltenberg for the initial support with the fungal trait data curation.

## Author contributions

A.L. and N.A.B conceived and designed the study. A.L. performed the investigation and developed the methodology. V.R. developed software, with contributions from A.L. and C.T. The following authors, A.L., C.T., A.M., and L.F. performed formal analysis, including data curation, statistical analysis, and visualization. N.A.B and A.L. acquired funding for this project. A.L. wrote the original draft, with contributions from A.M. and L.F. All the authors critically revised the manuscript for its intellectual content and approved the final version.

